# Variants in the *FTO* and *CDKAL1* loci have recessive effects on risk of obesity and type 2 diabetes respectively

**DOI:** 10.1101/027490

**Authors:** Andrew R Wood, Jessica Tyrell, Robin Beaumont, Samuel E. Jones, Marcus A. Tuke, Katherine S. Ruth, The GIANT consortium, Hanieh Yaghootkar, Rachel Freathy, Anna Murray, Timothy M. Frayling, Michael N. Weedon

## Abstract

Genome-wide association studies have identified hundreds of common genetic variants associated with obesity and Type 2 diabetes. These studies have focused on additive association tests. Identifying deviations from additivity may provide new biological insights and explain some of the missing heritability for these diseases.

To identify non-additive associations we performed a genome-wide association study using a dominance deviation model for BMI, obesity and Type 2 diabetes (4,040 cases) in 120,286 individuals of British ancestry from the UK Biobank study.

Known obesity-associated variants in *FTO* showed strong evidence for deviation from additivity (*P*=3×10^−5^) through a recessive effect of the BMI-increasing allele. The average BMI of individuals carrying 0, 1 or 2 BMI-raising alleles was 27.27kg/m^2^ (95% CI:27.22-27.31), 27.54kg/m^2^ (95% CI:27.50-27.58), and 28.07kg/m^2^ (95% CI:28.0-28.14), respectively. A similar effect was observed in 105,643 individuals from the GIANT consortium (*P*=0.003; *P_meta-analysis_*=1×10^−7^). We also detected a recessive effect (*P*_domdev_=5×10^−4^) at *CDKAL1* for Type 2 diabetes risk. Homozygous risk allele carriers had an OR=1.48 (95% CI:1.32-1.65) in comparison to the heterozygous group that had an OR=1.06 (95% CI:0.99-1.14), a result consistent with a previous study. We did not identify any novel genome-wide associations.

In conclusion, although we find no evidence for widespread non-additive effects contributing to the genetic risk of obesity and Type 2 diabetes, we find robust examples of recessive effects at the *FTO* and *CDKAL1* loci.

## INTRODUCTION

Genome-wide association (GWA) studies have identified hundreds of variants associated with obesity and T2D (1-9). GWA studies of T2D and obesity have usually focused on testing additive models. An additive model assumes that the disease risk of heterozygous individuals is exactly halfway between those of the two homozygous groups. Non-additive effects include dominant and recessive effects. These effects are common in monogenic disorders, but there are only a few examples in common diseases and traits (10). For obesity and T2D, the strongest evidence for a non-additive effect is at the *CDKAL1* locus where one study demonstrated a recessive effect (11). The GIANT consortium tested 32 BMI-associated variants for deviations from the additive model but, overall, found no evidence for deviation from additivity in 105,643 individuals (5).

There are several reasons why it is important to test for non-additive associations between common genetic variants and T2D and obesity. First, a genome-wide approach that tests alternative models could identify new variants and candidate genes. Second, variants that operate through a predominately recessive or dominant model could provide insights into the mechanism of action of variation at the associated loci providing new insights into biology, and help guide functional studies. Third, the correct model of inheritance could explain more of the variation in the trait, and hence account for some of the “missing heritability”.

The UK Biobank provides an excellent opportunity to test for deviation from additivity in a single large cohort as genome-wide SNP and detailed phenotypic data is available in the initial release of data from over 120,000 British individuals (12). In this study we used the UK Biobank to perform genome-wide association tests for deviations from the additive model for BMI, Obesity and T2D.

## METHODS

### Samples

We used 120,286 individuals of British descent from the first UK Biobank genetic data release. Basic characteristics are given in **Supplementary Table 1**. British-descent was defined as individuals who both self identified as white British and were confirmed as ancestrally Caucasian using principal components analyses (http://biobank.ctsu.ox.ac.uk).

### Genotypes

We used directly genotyped data from the UK Biobank for genome-wide association analyses. We limited our analyses to directly typed variants and only used imputed data for variants known to be associated with BMI or T2D. Using the 120,286 individuals, SNPs were excluded if not in Hardy-Weinberg Equilibrium (HWE) (*P*<1×10^−6^) or had a minor allele frequency <1%. In addition, we excluded SNPs with an overall missing rate >1.5% across all UK Biobank participants. This resulted in 484,426 directly genotyped SNPs for analysis. For known BMI and T2D variants, we extracted genotypes from UK Biobank’s imputation dataset and converted to best-guess genotypes prior to association testing. Genotypes were excluded if the genotype probability < 0.9. SNPs were excluded if HWE *P*<1×10^−6^ or failed imputation quality control (imputation quality < 0.9)

### Selection of known variants

#### BMI and Obesity

We selected common genetic variants that were associated with BMI in the GIANT consortium (5). We limited the BMI SNPs to one per locus and those that were associated with BMI in the analysis of all European ancestry individuals. In total, 72 SNPs previously associated with BMI were analysed **(Supplementary Table 2)**.

#### Type 2 diabetes

We selected common genetic variants previously associated with T2D in the DIAGRAM consortium (3). Details of the 66 T2D SNPs are provided in **Supplementary Table 3**.

### Within British Principal Components Analysis

The UK Biobank study identified 120,286 individuals who were both self-identified as white British and confirmed as ancestrally Caucasian using genetics and principal components analyses. The 120,286 individuals also excluded third degree or closer relatives. We performed an additional round of principal components analysis (PCA) on these 120,286 UK Biobank participants. We selected 95,535 independent SNPs (pairwise *r*^2^ >0.1) directly genotyped with a minor allele frequency (MAF) ≥ 2.5% and missingness <1.5% across all UK Biobank participants with genetic data available at the time of this study (*n*=152,732), and with HWE *P*>1×10^−6^ within the white British participants. Principal components were subsequently generated using FlashPCA (13).

### Phenotypes

#### BMI

The UK Biobank provides two measures of BMI – one calculated from weight (kg)/height^2^ and one using electrical impedance. We excluded individuals with differences >4.56 SDs between impedance and normal BMI measures where both variables were available. If only one measure of BMI was available this was used. We corrected BMI by regressing age, sex, study centre, and the first 5 within-British principal components and taking residual values. We then inverse normalised the residuals and used this analysis as our primary result. We also ran the genome-wide tests without inverse normalisation of BMI to avoid any potential attenuation of power by truncating the naturally right-hand skewed distribution of BMI. A total of 119,688 white-British individuals with BMI and genetic data were available.

#### “Obese” and “morbidly obese” categorical variables

Individuals were classified as “obese” if their BMI ranged from 30 to 34.9kg/m^2^, (N=29,925) and “severely obese” if their BMI exceeded this range (≥35kg/m^2^) (N=2,389). Controls for both were defined as those with a BMI <25kg/m^2^.

#### Type 2 diabetes

Individuals were defined as having T2D if they reported either T2D or generic diabetes at the interview stage of the UK Biobank study. Individuals were excluded if they reported insulin use within the first year of diagnosis. Individuals reportedly diagnosed under the age of 35 years or with no known age of diagnosis were excluded, to limit the numbers of individuals with slow-progressing autoimmune diabetes or monogenic forms. Individuals diagnosed with diabetes within the last year of this study were also excluded as we were unable to determine whether they were using insulin within this time frame. A total of 4,040 cases and 113,735 controls within the white British subset of UK Biobank were identified with genetic data available.

### Genome-wide association analysis

We adjusted for genotyping chip at run-time for the analyses of additive, dominance deviation, and recessive models for both BMI and T2D. Standard linear and logistic regression methods were applied to BMI and T2D, respectively. Logistic regression models included age, sex, study centre, and the first 5 within-British principal components as additional covariates.

### Genome-wide detection of deviation from additivity

We used a dominance deviation test to test for deviation for additivity. We performed a regression analysis against our trait of interest with a term representing the additive model for genotypes (coded 0,1,2 for homozygote, heterozygous, alternate homozygote group) and a term for dominance deviation (coded 0,1,0).

## RESULTS

### Genome-wide association study for deviation from additivity for BMI

We did not observe evidence for deviation from additivity at any SNP for BMI at genome-wide significance, with BMI either under an inverse-normalised or natural scale. **Supplementary Figure 1** presents the genome-wide QQ plots for all directly genotyped variants for the dominance deviation test on the inverse-normalised scale. **Supplementary Table 4** presents the top genome-wide association signals for deviation from additivity (*P_DOMDEV_*<1×10^−4^).

### Alleles at the FTO locus have a partially recessive effect on BMI and obesity status

A variant, rs9940128, representing the known BMI signal at the *FTO* locus had the 4^th^ strongest association with deviation from additivity (*P_DOMDEV_*=3×10^−6^). This variant is in very strong linkage disequilibrium (LD) (*r*^2^ = 0.92, *D’* > 0.99) with rs1421085 recently identified as the best candidate causal variant at *FTO* (2, 14). We used imputed rs1421085 genotypes to show that this SNP similarly deviated from a standard additive model (*P_DOMDEV_*=3×10^−5^) as shown in **Table 1** and **Supplementary Table 5**.

**Table 1.** Summary statistics for the most strongly associated directly genotyped SNP (rs9940128) and previously reported index SNP (rs1421085) at the *FTO* locus (*r*^2^ = 0.92) with evidence of departure from the additive model. Effect sizes are derived from BMI after inverse-normalisation of covariate adjusted residuals.

| SNP | Effect/<br>Other Allele | <u>Additive Effects</u> |  |  | <u>Deviation from Additivity</u> |  |  |
| --- | --- | --- | --- | --- | --- | --- | --- |
| | | $\beta$ | s.e. | *P* | $\beta$ | s.e. | *P* |
| rs1421085 | C/T | 0.076 | 0.004 | 2E-75 | -0.025 | 0.006 | 3E-05 |
| rs9940128 | A/G | 0.073 | 0.004 | 2E-69 | -0.027 | 0.006 | 3E-06 |

Homozygous carriers of the BMI raising allele had an average BMI of 28.07 kg/m^2^ (95% CI: 28.0-28.14 kg/m^2^), heterozygotes had an average BMI of 27.54 kg/m^2^; (95% CI: 27.50-27.58 kg/m^2^) and homozygous carriers of the BMI lowering allele an average BMI of 27.27 kg/m^2^; 95% CI: 27.22-27.31 kg/m^2^) (**Fig. 1A** and **Table 2**). While heterozygous carriers were still on average more overweight than the common allele homozygote group, the difference (0.27 kg/m^2^) was approximately half that between the heterozygote and minor allele homozygote groups (0.53 kg/m^2^). Accounting for this partially recessive effect only resulted in a small increase in variance of BMI explained (an additional 0.01%).

**Figure 1.**
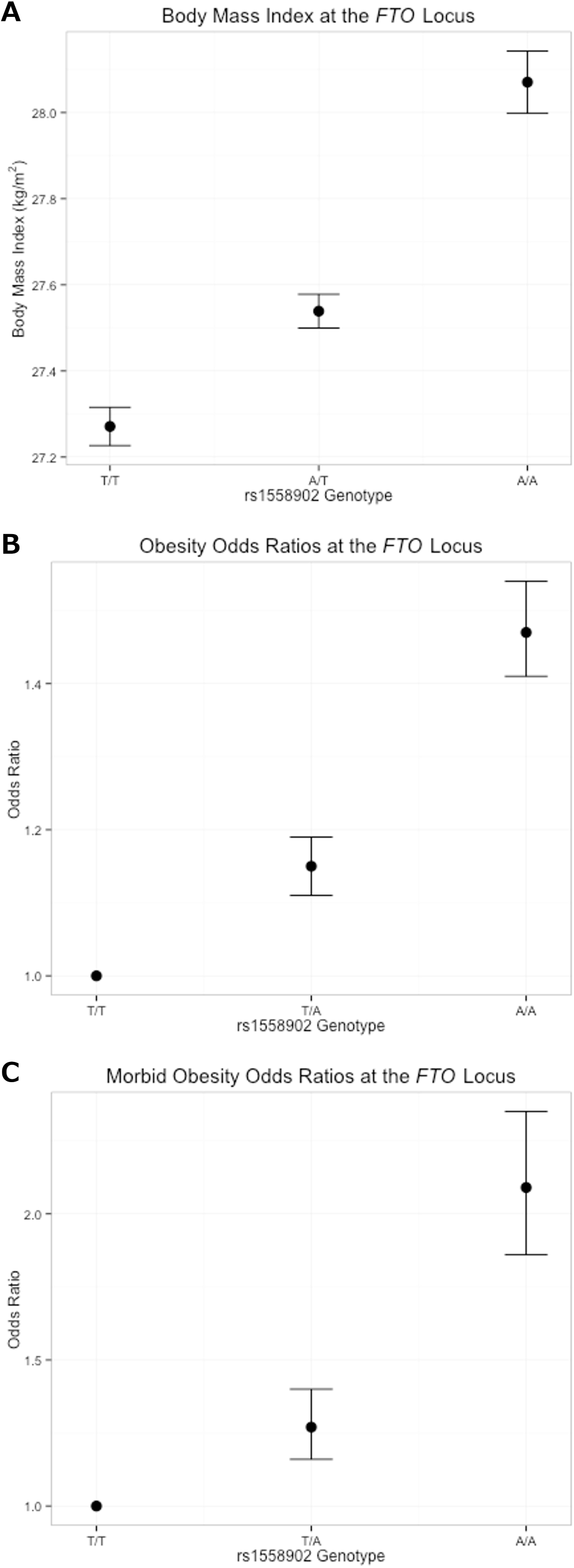
Average BMI and obesity odds ratios with 95% confidence intervals for carriers of the BMI raising allele at the *FTO* locus represented by rs1421085. A: Average BMI within each of the three genotype classes. B: Obesity risk for the heterozygous and homozygous carriers of the BMI increasing allele. C: Morbid obesity risk for the heterozygous and homozygous carriers of the BMI increasing allele.

**Table 2.**
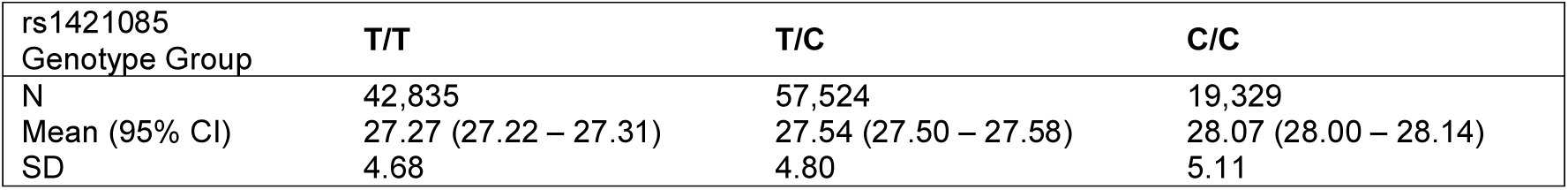
BMI values by genotype group at the *FTO* locus (rs1421085) for 119,688 white British individuals available in the UK Biobank. Allele ‘C’ is the BMI raising allele. Units are in kg/m^2^. 95% CI = 95% confidence interval, SD = standard deviation.

| rs1421085<br>Genotype Group | T/T | T/C | C/C |
| --- | --- | --- | --- |
| N | 42,835 | 57,524 | 19,329 |
| Mean (95% CI) | 27.27 (27.22 – 27.31) | 27.54 (27.50 – 27.58) | 28.07 (28.00 – 28.14) |
| SD | 4.68 | 4.80 | 5.11 |

The *FTO* locus also showed a similar pattern of deviation from additivity in case/control analyses of obesity (*P_DOMDEV_*=0.001) and morbid obesity (*P_DOMDEV_*=0.003) (**Fig. 1B and C**, and **Table 3**). The OR was stronger in homozygous carriers of the risk allele than expected under an additive model. The observed OR for obese heterozygous carriers of the BMI raising allele was 1.15 (95% CI: 1.12-1.19). Under the additive model we would expect an OR~1.32 for obese homozygous carriers yet the observed OR was 1.48 (95% CI:1.41-1.55). Similarly, the OR observed in the morbidly obese heterozygote and homozygous carriers was 1.28 (95% CI: 1.16-1.41) and 2.10 (95% CI: 1.87-2.35), respectively, whereas under the additive model we would expect an OR ~1.64 for the homozygous carriers.

**Table 3.** Odds ratios for “obese” and “morbidly obese” classifications by genotype group at the *FTO* locus (rs1421085). Allele “C” is the risk increasing allele. OR = odds ratio, 95% CI = 95% confidence interval.

| Classification | <u>Additive</u><br>OR<br>(95% CI) | *P* | <u>T/T vs. T/C</u><br>OR<br>(95% CI) | *P* | <u>T/C vs. C/C</u><br>OR<br>(95% CI) | *P* | <u>T/T vs. C/C</u><br>OR<br>(95% CI) | *P* |
| --- | --- | --- | --- | --- | --- | --- | --- | --- |
| Obese | 1.20<br>(1.18-1.23) | 3E-60 | 1.15<br>(1.12–1.19) | 1E-16 | 1.28<br>(1.22–1.34) | 4E-28 | 1.48<br>(1.41–1.55) | 5E-62 |
| Morbidly Obese | 1.44<br>(1.35-1.52) | 7E-33 | 1.28<br>(1.16-1.41) | 8E-07 | 1.65<br>(1.48-1.83) | 4E-20 | 2.10<br>(1.87-2.35) | 6E-36 |

### The partially recessive effect at FTO is present in 105,643 individuals from the GIANT consortium

The GIANT consortium previously tested for deviations from additivity for 32 known BMI variants using 105,643 individuals (5). There was no overall evidence for deviations from additivity at these known loci; however, *FTO* did have a similar partial recessive effect in this independent dataset represented by rs1558902 - a proxy of rs1421085 (*r*^2^ > 0.99, *D’* = 1) (*P_GIANT_DOMDEV_* = 0.003; *β* = −0.019; 95% CI: −0.031,-0.008). The negative direction of effect for the heterozygous group in comparison to the two homozygous groups combined was consistent with that observed in UK Biobank (**Table 1**) and indicative of a recessive effect for the BMI increasing allele. Meta-analysing the studies strengthened the evidence of deviation from additivity (N=225,143; *P_META-ANALYSIS_=*1×10^−7^).

### No evidence for non-additive effects at other known BMI variants

There was no evidence of deviation from additive effects for the remaining 71 BMI variants (**Supplementary Table 5**). Based on the 72 BMI variants we also showed that using an inverse-normalised distribution of BMI produced very similar results as when using BMI on its naturally, skewed, scale (**Supplementary Figure 2**).

### Genome-wide association study for deviation from additivity for T2D

We did not identify any variants deviating from additivity for T2D at genome-wide significance (all variants *P_DOMDEV_*<1×10^−5^; **Supplementary Figure 3**).

**Supplementary Table 6** presents the top signals for dominance deviation (*P_DOMDEV_*<1×10^−4^).

### Alleles at the CDKAL1 locus have a recessive effect for T2D

We found evidence of deviation from additivity for the SNP rs7756992 at the *CDKAL1* locus for T2D status (*P_DOMDEV_*=5×10^−4^) (**Fig 2**. and **Table 4**). The genotype OR was stronger in homozygous carriers for the risk allele than expected under an additive model (**Table 5**). The observed OR within the heterozygous carriers of the risk-increasing allele was 1.06 (95% CI 0.99 to 1.14) (*P*=0.08) which is smaller than the expected OR of ~1.22 for heterozygous carriers under an additive model. This finding is consistent with a previous study by deCODE that showed evidence of this SNP having a similar pattern of association with T2D risk (11). We found no evidence for deviation from additivity for any of the remaining known T2D variants.

**Figure 2.**
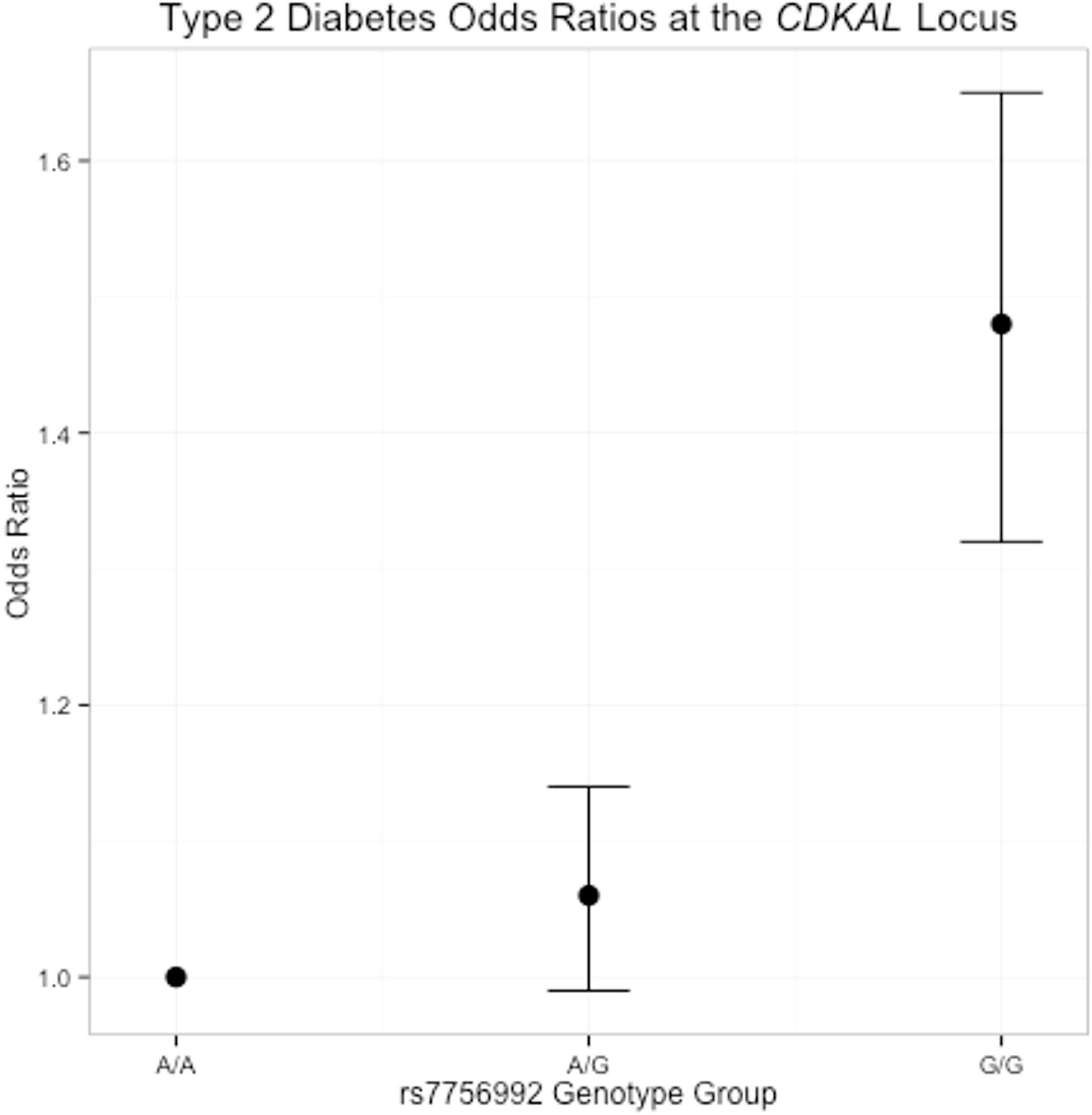
Odds ratios and 95% confidence intervals for the heterozygous and homozygous carriers of the *CDKAL1* T2D risk allele against the reference non-risk allele homozygous group.

**Table 4.** Summary statistics for rs7756992 at the *CDKAL1* locus showing evidence of deviation from additivity. The effect allele is the allele observed risk-increasing raising allele under the additive test.

| SNP | Locus | Risk/<br>Other Allele | <u>Deviation from Additivity</u> |  |  |
| --- | --- | --- | --- | --- | --- |
|  |  |  | *Odds Ratio* | *s.e.* | *P* |
| rs7756992 | *CDKAL1* | G/A | 0.87 | 0.038 | 5E-04 |

**Table 5.** T2D odds ratios by genotype groups at the *CDKAL1* locus (rs7756992) for 119,500 white British individuals available in the UK Biobank. Allele ‘G’ is the T2D risk increasing allele. OR = odds ratio, 95% CI = 95% confidence interval.

| <u>Allelic</u> |  | <u>A/A vs A/G</u> |  | <u>A/G vs G/G</u> |  | <u>A/A vs G/G</u> |  |
| --- | --- | --- | --- | --- | --- | --- | --- |
| OR<br>(95% CI) | *P* | OR<br>(95% CI) | *P* | OR<br>(95% CI) | *P* | OR<br>(95% CI) | *P* |
| 1.15<br>(1.10 – 1.21) | 1E-08 | 1.06<br>(0.99 – 1.14) | 0.077 | 1.39<br>(1.24 – 1.56) | 1E-08 | 1.48<br>(1.32 – 1.65) | 6E-12 |

## DISCUSSION

Our analyses of 120,286 UK Biobank individuals suggest that most genetic variants associated with BMI and T2D operate through a per-allele additive effect. Our findings suggest that dominant and recessive effects at common variants have a minimal role in explaining variation in BMI and risk of obesity and T2D. Our results are consistent with a previous smaller study of 6,715 individuals that concluded that deviations from additivity contribute little to missing heritability for a wide range of traits (15). There were exceptions for the *FTO*-BMI association and the *CDKAL1-*T2D association.

The 16% of individuals carrying two copies of the BMI raising allele at the *FTO* locus had more than twice the expected BMI difference compared to individuals carrying no BMI-raising alleles than would have been expected under a purely additive model. Assuming an average height in males of 1.78cm, it is equivalent to homozygous carriers of the BMI increasing allele being 2.53kg heavier than homozygous carriers of the opposite allele, whereas heterozygous carriers would be only 0.86kg heavier. Previous studies have shown that the vast majority of this increased weight is fat mass (1). The results are also consistent with a study of the *FTO* variant in polycystic ovarian syndrome (16). For T2D, we found evidence of a recessive effect at the *CDKAL1* locus. The evidence that heterozygous carriers of the risk allele were at increased risk of T2D was minimal and combined with previous data from the deCODE study suggests the true biological effect at this locus is recessive. Although accounting for non-additive effects at these loci only explained a small amount of additional variation in risk of obesity and T2D, understanding why these associations demonstrate non-additivity may provide new insights into biological mechanisms at these loci.

A strength of our study is that we used a single large relatively homogeneous dataset with full access to individual level genotype and phenotype data. We had >80% power to detect dominance deviation from additivity explaining 0.04% of the phenotypic variance at *P*=5×10^−8^. This is equivalent to being able to detect a purely recessive effect of 0.4 kg/m^2^ for a BMI allele with a frequency of 0.25, for example. Our analyses show how single large studies such as the UK Biobank will provide added value to existing meta-analyses approaches in GWA studies.

Our analyses have some limitations. We only analysed directly genotyped variants, and our statistical power to detect deviations from additivity may have been reduced if variants we analysed are imperfect markers for causal alleles. Non-biological explanations for the non-additive effects include “haplotype effects” due to LD with other causal alleles. In such situations, alleles of SNPs showing evidence of non-additivity are partially correlated with a much stronger causal SNP with an additive effect (17, 18). This is unlikely to be the case at the *FTO* or *CDKAL1* loci. These loci have been studied extensively through re-sequencing and fine mapping efforts and no substantially stronger individual variants have been identified, and for *FTO,* rs1421085 was recently proposed as the most likely causal variant (14).

The detection of non-additive genetic effects for BMI is potentially complicated by the skewed distribution of BMI. Effects that seem recessive could be artefacts of the skewed nature of the BMI distribution as variation in BMI is wider towards the more overweight end of the distribution. To limit this effect we inverse-normalised BMI and performed additional sensitivity analyses (including “robust regression” - an alternative to “least squares regression”) that accounts for different variances of a trait which may be the case for *FTO* (19) data not shown) to limit the influence of the skewed distribution, and found no evidence of BMI increasing alleles being more likely to have recessive effects than BMI lowering alleles (see **Supplementary Table 5**, and **Supplementary Figures 4** and **5**). Alternatively artificially truncating the BMI distribution into a normal distribution could reduce power to detect recessive effects of BMI-increasing alleles. However, we saw very little reduction in statistical confidence of known BMI associations when using the inverse-normalised scale compared to the natural BMI scale.

In conclusion, we have performed tests of deviation from additivity for BMI, obesity and T2D. Overall, there is little evidence of dominant and recessive effects. However, we find replicable examples of non-additive effects at *FTO* on BMI and obesity, and *CDKAL1* on T2D risk. Recessive effects have implications for the mechanism of action of these loci but do not explain appreciably more of the “missing heritability”.

## ACKNOWLEDGEMENTS

This research has been conducted using the UK Biobank Resource.

**Funding Information**. A.R.W. and T.M.F. are supported by the European Research Council grant: 323195: SZ-245 50371-GLUCOSEGENES-FP7-IDEAS-ERC. R.M.F. is a Sir Henry Dale Fellow (Wellcome Trust and Royal Society grant: 104150/Z/14/Z). R.B. is funded by the Wellcome Trust and Royal Society grant: 104150/Z/14/Z. J.T. is funded by the ERDF and a Diabetes Research and Wellness Foundation Fellowship. S.E.J. is funded by the Medical Research Council (grant: MR/M005070/1) M.A.T., M.N.W. and A.M. are supported by the Wellcome Trust Institutional Strategic Support Award (WT097835MF). H.Y. is funded by the European Research Council award (323195). The funders had no influence on study design, data collection and analysis, decision to publish, or preparation of the manuscript.

**Author Contributions**. A.R.W, T.M.F, M.N.W designed the study. A.R.W, T.M.F, M.N.W wrote the manuscript. A.R.W., J.T., R.B., S.E.J, M.A.T, K.S.R., H.Y., R.F., A.M., M.N.W performed data processing and statistical analyses.

M.N.W. is the guarantor of this work and, as such, had full access to all the data in the study and takes responsibility for the integrity of the data and the accuracy of the data analysis.

**Duality of Interest**. No potential conflicts of interest relevant to this article were reported.

